# A novel test for detecting gene-gene interactions in trio studies

**DOI:** 10.1101/021469

**Authors:** Brunilda Balliu, Noah Zaitlen

## Abstract

Epistasis plays a significant role in the genetic architecture of many complex phenotypes in model organisms. To date, there have been very few interactions replicated in human studies due in part to the multiple hypothesis burden implicit in genome-wide tests of epistasis. Therefore, it is of paramount importance to develop the most powerful tests possible for detecting interactions. In this work we develop a new gene-gene interaction test for use in trio studies called the trio correlation (TC) test. The TC test computes the expected joint distribution of marker pairs in offspring conditional on parental genotypes. This distribution is then incorporated into a standard one degree of freedom correlation test of interaction. We show via extensive simulations that our test substantially outperforms existing tests of interaction in trio studies. The gain in power under standard models of phenotype is large, with previous tests requiring more than twice the number of trios to obtain the power of our test. We also demonstrate a bias in a previous trio interaction test and identify its origin. We conclude that the TC test shows improved power to identify interactions in existing, as well as emerging, trio association studies. The method is publicly available at www.github.com/BrunildaBalliu/TrioEpi.

## INTRODUCTION

Genetic association studies, especially in humans, have focused primarily on marginal effects of genetic variants. While this approach has successfully identified thousands of variants associated with hundreds of complex human diseases (Hindorff *et al.* 2009), it ignores the role of epistasis in shaping phenotypes. Recent work in model organisms has shown that epistasis is a major contributor to broad sense heritability (Ayroles *et al.* 2009; Bloom *et al.* 2013; Ackermann and Beyer 2012) and interactions have been repeatedly posited as a key component of missing heritability in humans (Zuk *et al.* 2012; Gibson 2012). Furthermore, identification of epistatic interactions provides important insights into the functional organization of molecular pathways (Carlborg and Haley 2004; Brem *et al.* 2005; Cordell 2009; Ayroles *et al.* 2009; Ma *et al.* 2012; Ma *et al.* 2012; Ma *et al.* 2013).

One of the major obstacles in the identification of interactions in genetic association studies is the multiple hypothesis correction penalty induced by the examination of millions of pairs of SNPs. Therefore, it is of fundamental importance to develop the most powerful possible test statistic when searching for interactions. In this work we are concerned with tests for epistasis in trio studies in which mother-father-offspring trios are genotyped and the offspring is a carrier for the disease of interest, or the phenotype of interest is fitness.

There currently exist three classes of test for interaction between pairs of markers in trio studies. First, case-only correlation tests have been proposed where the null hypothesis is no correlation between genotypes at the two loci. While these can easily be performed via *χ*^2^ tests of independence between genotypes, they fail to leverage the information available from the parents in the trios and are susceptible to inflation from population structure. Second, pseudo-controls can be created via the non-transmitted parental alleles and used as matched controls in a conditional logistic regression framework (Cordell 2002; Cordell *et al.* 2004; Schwender *et al.* 2013). These tests account for population stratification that can induce long range LD between marker pairs and bias the case-only correlation tests. Third, and most recently, Ackermann and Beyer (2012) proposed the Imbalanced Allele Pair frequencies (ImAP) test. Their insight was that the expected counts of offspring alleles at a pair of SNPs could be computed conditional on parental genotypes. They incorporated this into a four degrees of freedom (d.f.) correlation test and used permutations to determine the null distribution (Ackermann and Beyer 2012).

The primary contributions of this work are two-fold. First, we show that in the presence of marginal effects but absence of interaction effects between pairs of markers, the ImAP test is biased. We identify that the normalization procedure used in ImAP is the source of this bias. Second, we develop a new interaction test, called the trio correlation test (TC) for use in trio studies. Using the insight of the ImAP approach, we begin by computing the expected distribution of the offspring’s genotype conditional on the parental genotypes and use this distribution to build a correlation test with one d.f. The TC has several advantages over previous approaches. First, it has one instead of four d.f. needed for ImAP. Second, it does not require expensive permutation tests to compute p-values. Third, we show via extensive simulations that our test is the most powerful among previously proposed tests under standard disease models. Indeed, in simulations where SNPs contained marginal and interaction effects, our average test statistic (16.4) was more than double the standard correlation based test statistic (7.8).

The rest of the paper is organized as follows. In Section 2, we introduce the existing tests and our proposed TC test. In Section 3, we evaluate the finite sample performance of the existing and proposed tests using an extensive simulation study. We close with a discussion in Section 4.

## METHODS

Consider a trio study in which *n* mother-father-offspring trios are genotyped, and the offspring are carriers for the disease of interest, or the phenotype of interest is fitness. In this section we present existing tests for detecting interaction between pairs of markers as well as our novel TC approach.

### The Standard Independence (SI) Test

Consider a pair of bi-allelic markers with possible genotypes *g*_1_, *g*_2_ ∈ {0, 1, 2}. Let *O*(*g*_1_, *g*_2_) the observed counts for the nine possible genotype combinations at these two markers in the offspring. Further, let *O*(*g*_1_) = *g*2 *O*(*g*_1_, *g*_2_) and *O*(*g*_2_) = *g*1 *O*(*g*_1_, *g*_2_) the observed marginal counts of the three possible genotypes at each marker. Last, let *E*(*g*_1_, *g*_2_) the expected counts of all nine possible genotype combinations at the two markers, as computed from the products of the observed marginal counts at each marker. That is *E*(*g*_1_, *g*_2_) = *O*(*g*_1_) *× O*(*g*_2_)*/n*.

A *χ*^2^ test statistic can be obtained by first calculating the squared difference of observed and expected counts for each genotype combination of the two markers divided by the corresponding expected counts. The final score for a marker pair is the sum of these values over all nine possible genotype combinations,

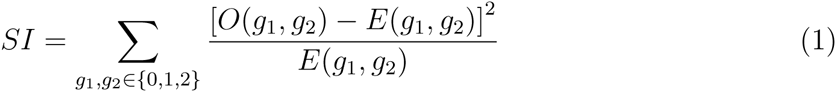

Under the null hypothesis of marker independence, *SI* is assumed to be asymptotically 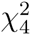 distributed.

### The Imbalanced Allele Pair Frequencies (ImAP) Test

Ackermann and Beyer (2012) proposed to calculate the expected counts in the children using the parental genotypes and the laws of Mendelian inheritance. Under Mendelian segregation, the offspring inherits alleles randomly from its parents and the expected genotype of each marker can be derived from the genotypes of the parents. The resulting probabilities for all possible parental genotype combinations are shown in Table 1.

**Table 1:**
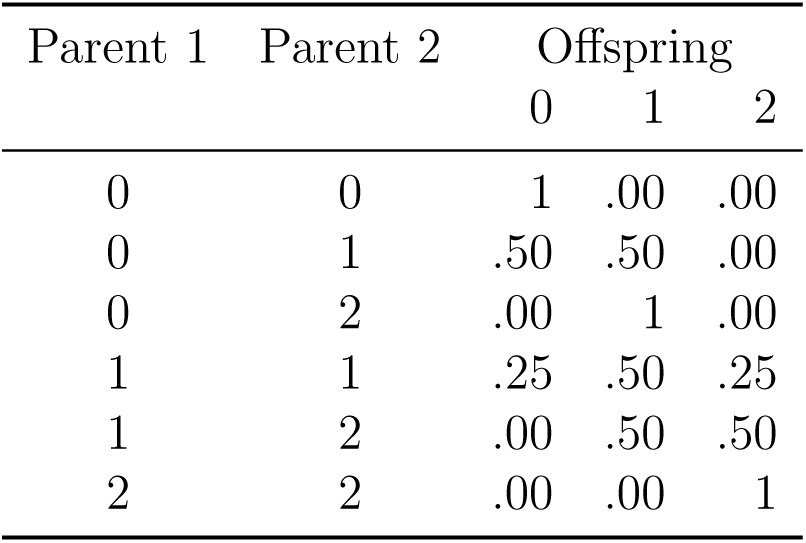
Expected genotype probabilities of an offspring given the parental genotypes according to Mendelian inheritance law.

| Parent 1 | Parent 2 | Offspring |  |  |
| --- | --- | --- | --- | --- |
|  |  | 0 | 1 | 2 |
| 0 | 0 | 1 | .00 | .00 |
| 0 | 1 | .50 | .50 | .00 |
| 0 | 2 | .00 | 1 | .00 |
| 1 | 1 | .25 | .50 | .25 |
| 1 | 2 | .00 | .50 | .50 |
| 2 | 2 | .00 | .00 | 1 |

Let 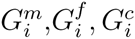 the genotypes of the mother, father and child of trio *i* at a marker.

Moreover, let

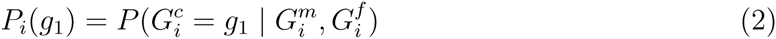

the probability of the offspring having genotype *g*_1_ conditional on the parental genotypes, as calculated using the probabilities in Table 1. For example, if 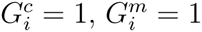 and 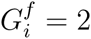then *P*_*i*_(0) =.25, *P*_*i*_(1) = .5, *P*_*i*_(2) = .25.

If the offspring are selected based on a phenotypic designation such as disease status, then a SNP increasing risk for the disease will be non-randomly inherited by the offspring. In order to correct for such main effects Ackermann and Beyer (2012) proposed to multiply each offspring’s expected genotype by the ratio of the sample-wide observed and expected counts for the corresponding marker, that is

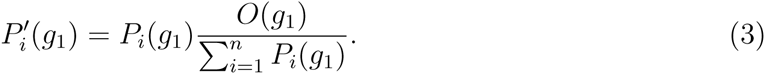

The above computation of 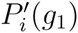 does not guarantee that 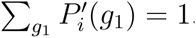 Ackermann and Beyer (2012) proposed to use the following normalization:

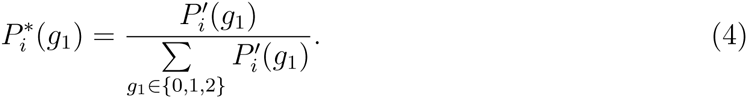

Subsequently, the expected counts of each genotype combination using the adjusted for main effects and normalized genotype counts can be calculated as

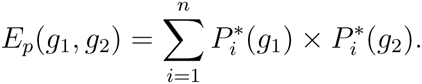

The corresponding *ImAP* statistic is given as,

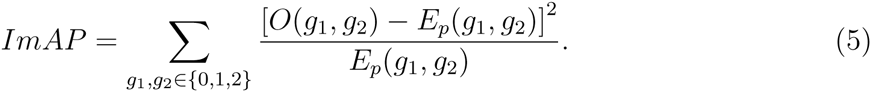

Because this *χ*^2^-like test statistic is not properly calibrated under the null hypothesis, Ackermann and Beyer (2012) assess the significance via a permutation approach. *ImAP* is nearly *χ*_4_^2^ distributed and we use this approximation to compute p-values in addition to the permutation approach. However, we show in Section 3 that in the presence of main effects the ImAP test statistic is inflated, even when the permutation approach is used to compute the p-values. The source of this inflation is the normalization step (4) (see Supplementary Text).

### The Standard Correlation (SC) Test

Let 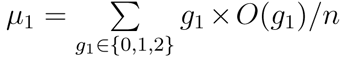 the expected value of the genotype at a marker and let 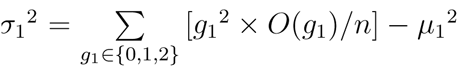 the corresponding variance. Moreover, let /inline/the covariance of 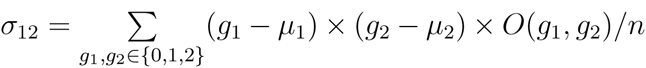 genotypes at the two markers and 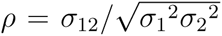their Pearson’s correlation coefficient.

The test statistic is given as

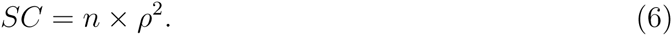

Under the null hypothesis, *SC* is assumed to be asymptotically 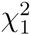 distributed.

### The Trio Correlation (TC) Test

Let 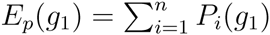 the expected genotype counts of genotypes at each of the two markers in the pair, computed based on the un-adjusted for main effects and un-normalized conditional genotype counts of the offspring in (2). In order to extend the standard correlation test based on the Pearson’s correlation coefficient such that information from parental genotypes is incorporated, we propose to compute *μ*_1_ and *μ*_2_ from the expected counts *E*_*p*_(*g*_1_) and *E*_*p*_(*g*_2_), rather than the observed genotype counts *O*(*g*_1_) and *O*(*g*_2_).

Let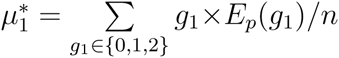the new expected value and 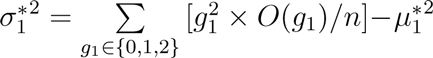the new variance. Moreover, let 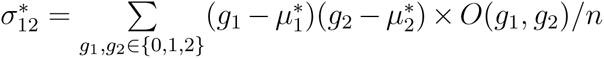the new covariance and 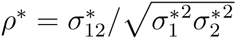the new correlation coefficient. Then the TC test statistic is given as

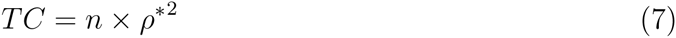

Under the null hypothesis, *T C* is assumed to be asymptotically 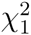 distributed. Unlike the *ImAP* test we do not utilize the normalized conditional probabilities and so we do not suffer from this source of inflation.

### Conditional Logistic Regression with Pseudo-controls (CLRPC)

In addition to the correlation and independence based tests described above, tests based on CLRPC have been proposed for detecting epistasis in trio studies. Briefly, fifteen pseudo-controls are constructed via the Mendelian genotype realizations, given the parents’ genotypes at the two loci, which are then used as matched controls in a CLRPC. Here we consider two types of tests based on CLRPC. First, interactions might be tested with a likelihood ratio test such as the one proposed by Cordell (2002). In this case, two conditional logistic regression models are fitted to the cases and the respective matched pseudo-controls, one consisting of two coding variables for each of the two SNPs, and the other additionally containing the four possible interactions of these variables. Then, p-values can be computed by approximation to a *χ*^2^ distribution with four d.f. We refer to this test as *CLRP C*_1_.

Alternatively, a more simple approach can be used in which interactions might be tested with a likelihood ratio test comparing a conditional logistic regression model containing one parameter for each SNP and one parameter for the interaction of these two SNPs with a model only consisting of the two parameters for the main effects of the SNPs, where a single genetic mode of inheritance assumed for each SNP (e.g. additive, dominant, or recessive). In this work we use additive marginal effects for both SNPs. The p-values can be computed by approximation to a *χ*^2^ distribution with one d.f. We refer to this test as *CLRP C*_2_.

In this work we use the R (R Core Team 2014) implementation of these tests available in the R package trio (Schwender *et al.* 2013) at http://cran.r-project.org.

## SIMULATION STUDY

### Data Generation

To evaluate the relative performance of the tests described in the previous section in terms of type I error rate and power, we performed a series of simulation studies under a liability threshold model of disease. Using random mating and Hardy-Weinberg Equilibrium assumptions we generated genotypes at two markers, each with a minor allele frequencies of 0.5, for a random sample of *N* trios, with *N* =10K, 50K and 100K. To generate *y*_*i*_, the liability of offspring *i*, we used the following regression model:

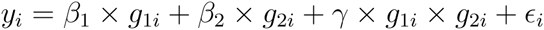

where *β*_1_ and *β*_2_ are the main effects of each marker and *γ* is their interaction effect, *∊*_*i*_ *∼ N* (0, *σ*^2^) a random error term with variance *σ*^2^ chosen such that both markers explain 10% of the heritability of the liability. This large effect size is used to test for inflation of interaction tests under main effects only models of phenotype.

In order to study the impact of ascertainment and disease prevalence in the performance of the tests, we selected from the initial random sample of *N* trios the 1,000 offspring with the highest phenotype values and their parents. The higher the value of *N*, the stronger the ascertainment is and the lower the disease prevalence is, since the sample of 1,000 trios represents a smaller fraction of the initial sample.

For each choice of parameters 10,000 data sets were generated and we applied the SI, ImAP, SC, TC, as well as the two CLRPC tests to each data set. We also examined the ImAP test without the normalization step (4), which we refer to as ImAP_2_. The *ImAP*_2_ test statistic is computed as the ImAP test statistic in (5) but the expected counts of each genotype combination are computed using the product of the adjusted for main effects only genotype counts *P*′, as opposed to the product of the adjusted for main effects and normalized counts *P ^*^* used for *ImAP*. The reason we show results for ImAP_2_ is to present the source of bias in the ImAP test, i.e. the normalization step in (4).

In addition, we also examined the ImAP test without the adjustment for main effects (3) and without the normalization step (4), which we refer to as ImAP_3_. Similarly to the *ImAP*_2_ test statistic, the *ImAP*_3_ is computed as the ImAP test statistic in (5) but now the expected counts of each genotype combination are computed using the product of *P*, i.e. the genotype counts without main effect adjustment or normalization. Results for ImAP_3_ are presented in order to study the impact of main effect adjustment when main effects are not present.

### Results on Type I Error Rate

We first examined the type I error rate performance of the tests under the null hypothesis of no main effects and no interaction effects (*β*_1_=*β*_2_=*γ*=0). The results in Table 2.a show that all tests with the exception of the ImAP tests are well calibrated, with an average *χ*^2^ test statistic equal to the d.f. of the test. We then examined the robustness of the tests when main effects, but no interaction effects exist (*β*_1_=*β*_2_=0.1 and *γ*=0). The results displayed in Table 2.b show that all tests suffered some degree of inflation. This inflation is driven by the fact that marginally associated SNPs will become correlated in ascertained data (Zaitlen *et al.* 2012; Zaitlen *et al.* 2012). Ascertainment based inflation will increase as the disease prevalence decreases.

**Table 2:** Mean test statistic with standard errors (se) of each test under the null hypothesis of no interaction effect, i.e. *γ* = 0. Each entry represents an average over 10,000 simulated data sets.

| Tests | d.f. | Mean Test Statistic (se) |  |  |
| --- | --- | --- | --- | --- |
| | | $N=10K$ | $N=50K$ | $N=100K$ |
| (a) $\beta_1 = \beta_2 = 0$ | | | | |
| $SI$ | 4 | 4.007 (.028) | 3.947 (.028) | 3.971 (.028) |
| $ImAP$ | 4 | 3.784 (.024) | 3.752 (.024) | 3.779 (.024) |
| $ImAP_2$ | 4 | 3.442 (.024) | 3.414 (.024) | 3.435 (.024) |
| $ImAP_3$ | 4 | 5.962 (.030) | 5.904 (.030) | 5.964 (.030) |
| $LRPC_1$ | 4 | 4.012 (.028) | 3.977 (.028) | 4.005 (.028) |
| $SC$ | 1 | 0.995 (.014) | 0.976 (.014) | 1.009 (.014) |
| $LRPC_2$ | 1 | 0.997 (.014) | 0.979 (.014) | 1.026 (.014) |
| $TC$ | 1 | 0.994 (.014) | 0.975 (.014) | 1.009 (.014) |
| (b) $\beta_1 = \beta_2 = .1$ | | | | |
| $SI$ | 4 | 5.637 (.038) | 5.533 (.037) | 5.485 (.036) |
| $ImAP$ | 4 | 22.183 (.036) | 35.700 (.042) | 41.011 (.044) |
| $ImAP_2$ | 4 | 4.509 (.030) | 4.615 (.031) | 4.615 (.030) |
| $ImAP_3$ | 4 | 84.660 (.129) | 154.454 (.185) | 185.087 (.207) |
| $LRPC_1$ | 4 | 5.279 (.036) | 5.280 (.036) | 5.248 (.036) |
| $SC$ | 1 | 2.747 (.030) | 2.642 (.029) | 2.628 (.029) |
| $LRPC_2$ | 1 | 2.224 (.026) | 2.188 (.026) | 2.174 (.026) |
| $TC$ | 1 | 0.958 (.013) | 2.415 (.027) | 3.949 (.036) |

Interestingly, the TC test is the least biased of these tests for the highest prevalence disease and the most biased for the lowest prevalence disease. This known source of bias is relatively minor compared to the genome-wide testing threshold and therefore unlikely to result in false-positive interaction results. However, the ImAP test is severely inflated under this marginal effects only model and is very likely to produce false positives. As noted in the previous section, this inflation is driven by the normalization procedure as evidenced by the fact that without normalization (ImAP_2_), the test is similar to the other tests considered.

It is well known that the *ImAP* tests are not *χ*^2^ distributed and so we computed the permutation-based p-values for the *ImAP*, *ImAP*_2_, and *ImAP*_3_ test statistics as described in Ackermann and Beyer (2012). The type I error rate for each of the three test based on the permutation p-values are listed in Table 3. Under the null hypothesis of no interaction and the absence of main effects, all three tests are well calibrated (Table 3.a). In the presence of main effects, *ImAP* and *ImAP*_3_ are very inflated. *ImAP*_3_ is inflated due to the presence of main effects. *ImAP*_2_ adjusts to some degree for the presence of main effects and has a much lower type I error rate. However, in the recommended *ImAP* test, the normalization step is performed which leads to an increased type I error rate even after performing permutations.

**Table 3:**
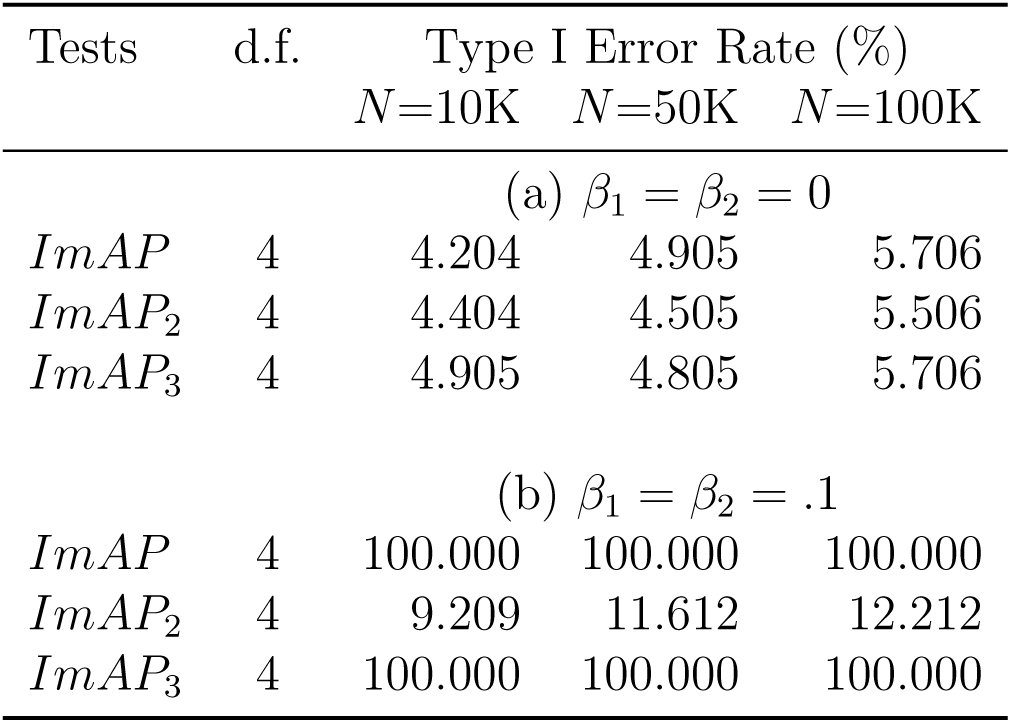
Permutation-based type I error rate for the ImAP, ImAP2 and ImAP3 tests under the null hypothesis of no interaction effect, i.e. *γ* = 0. Each entry is based on 1,000 permuted data sets.

Population structure can induce correlation between distant SNPs. In order to examine the effects of structure on the interaction tests we performed another a set of simulations in a pseudo-admixed population. We assumed minor allele frequencies of 0.5 and 0.1 in populations one and two respectively. For each mother and father we drew genome-wide ancestry uniformly on the interval [0.1,0.9]. We then drew local ancestry from a binomial according to the genome-wide ancestry. Finally, we drew genotypes based on the minor allele frequencies for each population conditional on the local ancestry at each SNP. The rest of the simulation proceeded as above.

The results of the pseudo-admixed population simulations under the null hypothesis of no interaction effect are presented in Table 4. As expected all interactions tests are inflated in structured populations. However, the CLRPC test is less inflated than all others and is therefore the recommended test when dealing with structured populations.

**Table 4:**
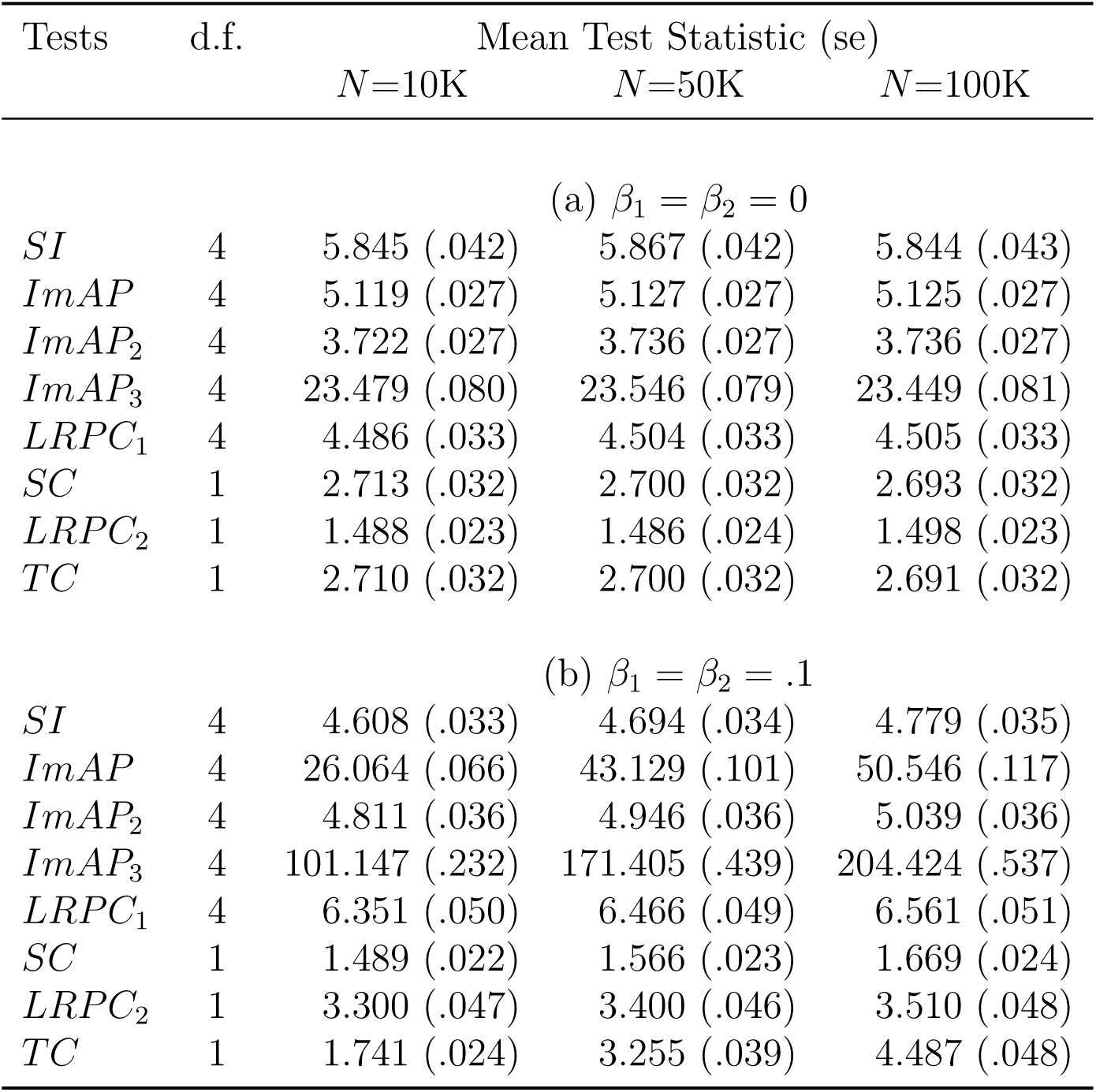
Mean test statistic with standard errors (se) of each test under the null hypothesis of no interaction effect, i.e. *γ* = 0, in the presence of admixture.

| Tests | d.f. | Mean Test Statistic (se) |  |  |
| --- | --- | --- | --- | --- |
| | | $N=10K$ | $N=50K$ | $N=100K$ |
| (a) $\beta_1 = \beta_2 = 0$ | | | | |
| $SI$ | 4 | 5.845 (.042) | 5.867 (.042) | 5.844 (.043) |
| $ImAP$ | 4 | 5.119 (.027) | 5.127 (.027) | 5.125 (.027) |
| $ImAP_2$ | 4 | 3.722 (.027) | 3.736 (.027) | 3.736 (.027) |
| $ImAP_3$ | 4 | 23.479 (.080) | 23.546 (.079) | 23.449 (.081) |
| $LRPC_1$ | 4 | 4.486 (.033) | 4.504 (.033) | 4.505 (.033) |
| $SC$ | 1 | 2.713 (.032) | 2.700 (.032) | 2.693 (.032) |
| $LRPC_2$ | 1 | 1.488 (.023) | 1.486 (.024) | 1.498 (.023) |
| $TC$ | 1 | 2.710 (.032) | 2.700 (.032) | 2.691 (.032) |
| (b) $\beta_1 = \beta_2 = .1$ | | | | |
| $SI$ | 4 | 4.608 (.033) | 4.694 (.034) | 4.779 (.035) |
| $ImAP$ | 4 | 26.064 (.066) | 43.129 (.101) | 50.546 (.117) |
| $ImAP_2$ | 4 | 4.811 (.036) | 4.946 (.036) | 5.039 (.036) |
| $ImAP_3$ | 4 | 101.147 (.232) | 171.405 (.439) | 204.424 (.537) |
| $LRPC_1$ | 4 | 6.351 (.050) | 6.466 (.049) | 6.561 (.051) |
| $SC$ | 1 | 1.489 (.022) | 1.566 (.023) | 1.669 (.024) |
| $LRPC_2$ | 1 | 3.300 (.047) | 3.400 (.046) | 3.510 (.048) |
| $TC$ | 1 | 1.741 (.024) | 3.255 (.039) | 4.487 (.048) |

### Results on Power

Tables 5 and 6 show the mean *χ*^2^ and power results respectively under the alternative hypothesis of interaction effect, i.e. *γ*=0.1, for both absence of main effects, i.e. *β*_1_=*β*_2_=0, and presence of main effects, i.e. *β*_1_=*β*_2_=.1. In order to compare the relative performance of the tests, we selected a significance threshold *α* such that the power of the SI test was 50%. For all tests power increases as disease prevalence decreases since a larger genetic burden is required to exceed the liability threshold.

**Table 5:**
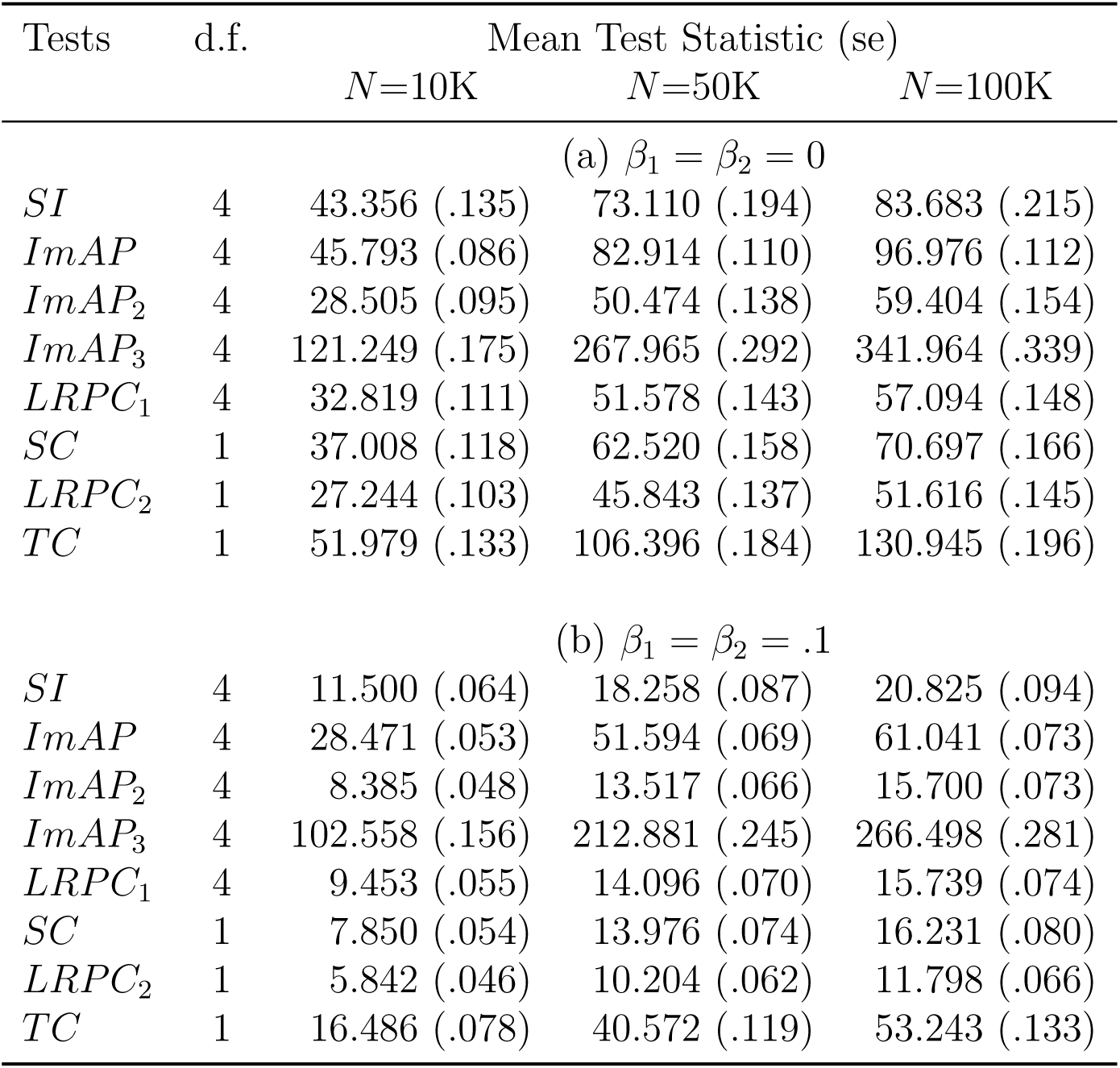
Mean test statistic with standard errors (se) of each test under the alternative hypothesis of interaction effect, *γ* = .1. Each entry represents an average over 10,000 simulated data sets.

| Tests | d.f. | Mean Test Statistic (se) |  |  |
| --- | --- | --- | --- | --- |
|  |  | N=10K | N=50K | N=100K |
| (a) $\beta_1 = \beta_2 = 0$ | | | | |
| *SI* | 4 | 43.356 (.135) | 73.110 (.194) | 83.683 (.215) |
| *ImAP* | 4 | 45.793 (.086) | 82.914 (.110) | 96.976 (.112) |
| *ImAP*<sub>2</sub> | 4 | 28.505 (.095) | 50.474 (.138) | 59.404 (.154) |
| *ImAP*<sub>3</sub> | 4 | 121.249 (.175) | 267.965 (.292) | 341.964 (.339) |
| *LRPC*<sub>1</sub> | 4 | 32.819 (.111) | 51.578 (.143) | 57.094 (.148) |
| *SC* | 1 | 37.008 (.118) | 62.520 (.158) | 70.697 (.166) |
| *LRPC*<sub>2</sub> | 1 | 27.244 (.103) | 45.843 (.137) | 51.616 (.145) |
| *TC* | 1 | 51.979 (.133) | 106.396 (.184) | 130.945 (.196) |
| (b) $\beta_1 = \beta_2 = .1$ | | | | |
| *SI* | 4 | 11.500 (.064) | 18.258 (.087) | 20.825 (.094) |
| *ImAP* | 4 | 28.471 (.053) | 51.594 (.069) | 61.041 (.073) |
| *ImAP*<sub>2</sub> | 4 | 8.385 (.048) | 13.517 (.066) | 15.700 (.073) |
| *ImAP*<sub>3</sub> | 4 | 102.558 (.156) | 212.881 (.245) | 266.498 (.281) |
| *LRPC*<sub>1</sub> | 4 | 9.453 (.055) | 14.096 (.070) | 15.739 (.074) |
| *SC* | 1 | 7.850 (.054) | 13.976 (.074) | 16.231 (.080) |
| *LRPC*<sub>2</sub> | 1 | 5.842 (.046) | 10.204 (.062) | 11.798 (.066) |
| *TC* | 1 | 16.486 (.078) | 40.572 (.119) | 53.243 (.133) |

**Table 6:** Power (%) of each test under the alternative hypothesis of interaction effect, *γ* = .1. Results are based on 10,000 simulated data sets.

| Tests | d.f. | Power (%) |  |  |
| --- | --- | --- | --- | --- |
| | | $N=10K$ | $N=50K$ | $N=100K$ |
| (a) $\beta_1 = \beta_2 = 0$ | | | | |
| $SI$ | 4 | 50.11 | 50.02 | 50.21 |
| $ImAP$ | 4 | 64.04 | 85.31 | 91.93 |
| $ImAP_2$ | 4 | 8.42 | 7.30 | 8.11 |
| $ImAP_3$ | 4 | 100.00 | 100.00 | 100.00 |
| $LRPC_1$ | 4 | 19.37 | 8.98 | 5.85 |
| $SC$ | 1 | 64.12 | 54.60 | 51.04 |
| $LRPC_2$ | 1 | 8.36 | 4.07 | 2.86 |
| $TC$ | 1 | 94.58 | 99.72 | 99.98 |
| (b) $\beta_1 = \beta_2 = .1$ | | | | |
| $SI$ | 4 | 50.01 | 49.30 | 48.96 |
| $ImAP$ | 4 | 100.00 | 100.00 | 100.00 |
| $ImAP_2$ | 4 | 27.99 | 25.76 | 25.42 |
| $ImAP_3$ | 4 | 100.00 | 100.00 | 100.00 |
| $LRPC_1$ | 4 | 36.57 | 29.43 | 26.50 |
| $SC$ | 1 | 68.73 | 68.59 | 66.72 |
| $LRPC_2$ | 1 | 14.72 | 13.69 | 12.00 |
| $TC$ | 1 | 96.57 | 99.93 | 100.00 |

The TC test is the most powerful test with a 40%, 70%, and 85% gain in test statistic relative to the SC test with a concomitant gain in power. The pseudo-control based interaction tests did not perform as well as the TC, SC and SI interaction tests. The SC test outperformed the SI tests due to the reduction in d.f and the additive generative model for the phenotype used in these simulations. The *ImAP* test is disqualified due to inflation introduced by the normalization. The ImAP_2_, which did not suffer this inflation, is underpowered relative to the other tests. By comparing ImAP_2_ to ImAP_3_ we can see that the reason ImAP_2_ is under-powered is mainly because of the ‘over-adjustment’ for main effects. ImAP_3_ is also disqualified as a proper test because it is severely inflated under the null hypothesis in the presence of main effects.

## DISCUSSION

In this work we develop the TC test, a new one d.f. statistical interaction test for use in trio studies that leverages information form the parental genotypes. We compare our test with existing tests for epistasis and show via simulations that the TC test properly controls the type I error in the absence of main effects and is similarly biased to other tests for low-prevalence diseases. Our test substantially outperforms all other tests of interaction in trio studies. The gain in power under standard models of phenotype is large, with the SC test requiring up to twice the number of trios to obtain the power of our test.

Ackermann and Beyer (2012) showed that in the absence of main effects the significance of the ImAP test statistic for each marker pair can be properly assessed via a permutation approach. We showed here that the permutation approach used to calibrate the ImAP test statistic in the absence of main effects will not address the inflation caused in the presence of main effects. We identify as a source of inflation the normalization step in (4), which alters the correlation structure of the expected genotype probabilities and introduces bias in the test statistic.

All methods considered here will be slightly inflated due to ascertainment for low prevalence diseases. However, this inflation is minimal and not enough to pass a genome-wide significance threshold. Population structure can induce long range LD inflating correlation based tests. Permutations based on pseudo-controls can account for structure (Epstein *et al.* 2012) if inflation is observed in the study. The logistic regression based pseudo-control method can directly account for population stratification and can flexibly model one d.f. up to four d.f. tests of interaction (Cordell *et al.* 2004). However, this test was the least powerful in our simulations and will lose power when there is assortative mating for the phenotype of interest (Klei *et al.* 2012).

In this work we used a liability threshold model (i.e. probit model) with and additive by additive interaction term to simulate disease phenotypes. Our previous work suggests that replacing this with a logit model will not have substantial effect on outcomes (Zaitlen *et al.* 2012). There exist a myriad of alternative models and assumptions each of which can alter the definition of interaction and the relative power of statistical tests (Cordell 2002; Hallgrímsdóttir and Yuster 2008). For example if the true model is four d.f.,e.g. contains additive by dominance effects, our test, which only models additive by additive effects, may no longer be the most powerful test. Because the ImAP test is inflated in the presence of main effects, a powerful and unbiased four d.f. test leveraging trios remains an open research question as does a one d.f. allelic test.

While we developed a correlation based test leveraging the parental genotypes, it also is possible to construct a likelihood based approach for interaction conditional on parental genotypes. This requires an assumption about the underlying disease model (e.g. logit/probit model). Such assumptions can lead to false conclusions of epistasis when the assumed model is incorrect (Clayton 2012). However, these tests can be more powerful when the model choices are similar to the real disease model.

Similar to commonly used tests for marginal effects, the interaction tests presented here are inappropriate for rare variants. In the marginal case it is recommended to use a Fisher’s Exact Test when the minor allele frequency is small. The definition of rare depends on the sample size of the study, as the number of observations of the rare allele is the quantity of interest. For interaction tests one must use a threshold on the product of the minor allele frequencies of pairs of markers.

There has been a renewed interest in trio cohorts with affected offspring for the purposes of identifying denovo mutations and parent of origin effects (O’Roak *et al.* 2011; Arjomandi *et al.* 2011; O’Roak *et al.* 2012; Neale *et al.* 2012; Sanders *et al.* 2012). While these collections have identified many denovo mutations, they have not yet been examined for the presence of interactions and our test is therefore of immediate benefit to these rapidly growing trio cohorts.

## ACKNOWLEDGEMENT

We thank Marit Ackermann and Andreas Beyer for helpful discussions. N.Z. was funded by 1K25HL121295-01A1.

## SUPPLEMENTARY TEXT

Let **p**_0_ = [*p*_1_(0), *…, p*_*n*_(0)], **p**_1_ = [*p*_1_(1), …, *p*_*n*_(1)], and **p**_2_ = [*p*_1_(2), …, *p*_*n*_(2)] vectors of the expected probability of genotypes 0, 1, and 2 for all offspring at a marker as calculated using the probabilities in Table 1 and let 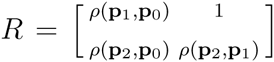 be the matrix of pairwise correlations between **p**_0_, **p**_1_, and **p**_2_. Similarly, let 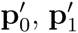 and 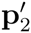 the vectors of the expected probabilities after main effects adjustment in (3), with *R*^*I*^ their correlation matrix; and 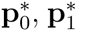 and 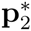 the vectors of expected probabilities after main effects adjustment and normalization in (4), with *R*^*′*^ their correlation matrix.

We showed in Section 3 that under the null hypothesis of no interaction effect but presence of main effects the *ImAP* tests statistic is inflated. We also showed that the ImAP permutation procedure does not address this inflation. Since *ImAP*_2_, the test without the normalization step, is not inflated under this scenario, we concluded that the normalization of the expected genotype probabilities in 4 is causing the inflation. We mention in the previous section that, in the presence of main effects, the normalization in (4) alters the correlation of the genotype probabilities used to compute the expected joint distribution of the markers in the cases but not in the pseudo-controls. To illustrate this, we compare the *R*, *R*^*′*^ and *R*^*\**^ matrices of the cases and pseudo-controls generated under the null hypothesis of no interaction and main effects, i.e. *γ* = *β*_1_ = *β*_2_ = 0, with the matrices for the cases and the pseudo-controls under the null hypothesis of no interaction effect, i.e. *γ* = 0, and presence of main effects, i.e. *β*_1_ = *β*_2_ = .1. These matrices are listed in Table 7.

**Table 7:** Correlation matrices of the expected genotype probabilities *R* (first column), the expected genotype probabilities after adjustment for main effects *R*^*′*^ (second column), and the expected genotype probabilities after adjustment for main effects and normalization *R*^*\**^(second column). Results are based on mean over 10,000 simulated data sets. PC, pseudo-controls.

| $R$ Cases | | | $R'$ Cases | | | $R^*$ Cases | | |
| --- | --- | --- | --- | --- | --- | --- | --- | --- |
|  | 0 | 1 |  | 0 | 1 |  | 0 | 1 |
| 1 | -0.446 |  | 1 | -0.446 |  | 1 | -0.447 |  |
| 2 | -0.601 | -0.447 | 2 | -0.601 | -0.447 | 2 | -0.600 | -0.447 |
| $R$ PC | | | $R'$ PC | | | $R^*$ PC | | |
|  | 0 | 1 |  | 0 | 1 |  | 0 | 1 |
| 1 | -0.445 |  | 1 | -0.445 |  | 1 | -0.445 |  |
| 2 | -0.600 | -0.449 | 2 | -0.600 | -0.449 | 2 | -0.599 | -0.449 |

(b) $\beta_1 = \beta_2 = .1$ , $\gamma = 0$
| $R$ Cases | | | $R'$ Cases | | | $R^*$ Cases | | |
| --- | --- | --- | --- | --- | --- | --- | --- | --- |
|  | 0 | 1 |  | 0 | 1 |  | 0 | 1 |
| 1 | -0.240 |  | 1 | -0.240 |  | 1 | -0.135 |  |
| 2 | -0.600 | -0.633 | 2 | -0.600 | -0.633 | 2 | -0.590 | -0.719 |
| $R$ PC | | | $R'$ PC | | | $R^*$ PC | | |
|  | 0 | 1 |  | 0 | 1 |  | 0 | 1 |
| 1 | -0.239 |  | 1 | -0.239 |  | 1 | -0.240 |  |
| 2 | -0.600 | -0.632 | 2 | -0.600 | -0.632 | 2 | -0.600 | -0.632 |

In the absence of main effects *R* = *R*^*I*^ and *R ≈ R^*^* (Table 7.a). On the other hand, in the presence of main effects the two matrices are similar for the pseudo-controls but are very different for the cases. The largest difference being between *ρ*_(_**p**_1_, **p**_0_) and 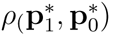 correlation of −0.240 before normalization and −0.135 after normalization, and between *ρ*_(_**p**_1_, **p**_2_) and 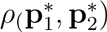 correlation of −0.633 before normalization and −0.719 after normalization.

